# Class-agnostic annotation of small RNAs balances sensitivity and specificity in diverse organisms

**DOI:** 10.1101/2025.01.25.634868

**Authors:** Nathan R. Johnson, Fabian Gonzalez-Toro, Barbara Bernal Gomez

**Affiliations:** Centro de Genómica y Bioinformática, Facultad de Ciencias, Ingeniería y Tecnología, Universidad Mayor, Santiago, Chile; Millennium Science Initiative - Millennium Institute for Integrative Biology (iBio), Santiago, Chile

## Abstract

Small RNAs (sRNAs) are important regulatory elements in eukaryotic organisms and comprise the functional elements of RNAi. Numerous classes of sRNAs have been annotated, however they vary greatly in their ease of annotation and compatibility with most annotators. Significant challenges exist for the annotation process, including variation in sRNA library quality, alignment depth, and poorly defined loci, collectively making this process difficult. Additionally, few annotators are fully agnostic to sRNA classes and may struggle identifying loci in less explored organisms (exceptional organisms, fungi). To address these problems, we present an integrated sRNA annotation suite, YASMA, which is specifically suited to finding reliable thresholds for locus annotation which balance sensitivity with specificity. By comparing YASMA with other annotators, we show that pipelines based on coverage-normalization methods have great advantages in balancing many metrics to produce a more reproducible annotation. We also demonstrate that YASMA produces more contiguous and representative loci, through the aggressive merging of similar adjacent expressed regions. Finally, we also show that the tool produces much more descriptive locus dimensions, a major advantage in species where sRNAs may be distinct or unique. Overall, we demonstrate substantial improvements in annotation accuracy, reproducibility, and description, particularly in non-model organisms and less-explored clades.

## Introduction

Small regulatory RNAs (sRNAs) are the functional elements behind RNA interference, a system of genetic regulation ubiquitous among eukaryotes. The core-machinery responsible for sRNAs are ancient and widely conserved **(Mukherjee et al., 2013; You et al., 2017)**. Processing follows the same general scheme: double-stranded RNA is processed by a type-III RNAse (Dicer or Dicer-like, here referred to collectively as DCR) and loaded into an Argonaute family protein which carries-out specific RNAi functions **(Zhan and Meyers, 2023; Torres-Martínez and Ruiz-Vázquez, 2017)**. Some sRNAs have been shown to be processed independent of DCR, such as piRNAs **(Vagin et al., 2006)** and some cases in fungi **(Lee et al., 2010)**. Within eukaryotic kingdoms, sRNA classes are mostly well conserved in processing and function, and some types are even identified across all eukaryotes **(Ghildiyal and Zamore, 2009; Axtell et al., 2011; Lee et al., 2010)**. Specific loci are also widely conserved, as seen with microRNAs (miRNAs) where orthologous loci are found in diverse organisms **(Cuperus et al., 2011)**. Despite these similarities, there are also many differences in the sRNA content and classes between disparate organisms and clades. This highlights the complexity and adaptability of sRNA these pathways across species.

In its most basic form, sRNA annotation provides a foundation for understanding the biology of sRNAs by classifying and clustering their expression into genic units. This usually involves identifying genomic loci, classification based on characteristics, and sometimes identification of homology and function. This process is used to help understand the sRNAs that exist in an organism and is often the first step to understanding their biology and function. Unlike in RNA-seq, where sequences come from the in vitro fragmentation of whole or larger transcripts, sRNA-genes are naturally fragmented by DCR **(Henderson et al., 2006)**. Moreover, sRNAs are short and sometimes sparsely aligned, making contig construction without a reference genome usually impossible. A prime example of this are miRNAs, which will by definition never produce a contiguous block **(Axtell and Meyers, 2018)**. Hairpin-precursors of miRNAs are sometimes detectable via directed amplification efforts in animals but are rarely, if-ever, found in plants **(Axtell et al., 2011)**. Other types of sRNA loci present even greater challenges due to their diverse precursors and less defined processing mechanisms. For instance, RNA-dependent RNA polymerase-derived siRNA loci in plants are shown to be produced by an assortment of intermediate-sized precursors, which are the products of many transcriptional initiations **(Blevins et al., 2015)**. While other mechanisms are known **(Peragine et al., 2004)**, the full details of transcription remain unknown for many sRNA-classes, particularly in less studied clades such as fungi.

Annotation of sRNAs is a common first-step in analysis. It follows that many annotators have been published, collectively with more than 5,500 citations to date (Fig. S1). However, these tools vary considerably in terms of annotation quality and focus. Most tools are exclusively focused on miRNAs, including the commonly used tools of the miRDeep-family **(Friedländer et al., 2012; Yang and Li, 2011)** and miRDeepfinder **(Xie et al., 2012)**. This preference is likely due to the relatively well-defined nature of miRNA loci **(Axtell and Meyers, 2018)** and their clear specific phenotypes **(Todesco et al., 2010)**. In contrast, less defined classes such as many siRNAs are more challenging to assess and are often ignored by annotators (Fig. S1). Nevertheless, these classes are crucial to understanding organisms that produce major portions of siRNAs (e.g., plants) **(Lunardon et al., 2020)**, piRNAs (e.g. some animals) **(Iwasaki et al., 2015)**, or those with hairpin RNAs that are less clearly defined (e.g. fungi) **(Lee et al., 2010; Johnson et al., 2022)**. Some tools do not perform *de novo* annotations, relying solely on previously published loci for their analysis (Fig. S1), which inherently limits highly-studied organisms and the limited published loci **(Griffiths-Jones et al., 2006)**. Other tools use genome-alignment-free approaches, dramatically narrowing the range of loci that can be discovered **(Vitsios et al., 2017; Yao et al., 2020)**. This leaves a vanishingly small set of annotators which might be capable of defining sRNAs in organisms with little or no definition of their sRNA classes **(Axtell, 2013)**.

Major hurdles persist in small RNA annotation. Library quality has a significant impact on problematic annotations **(Ludwig et al., 2017)**. Different organisms and clades exhibit distinct types of sRNAs. Most existing tools are primarily tested in a limited range of species, and often those with well-defined locus parameters. Critically, virtually no tools are designed for *de novo* sRNA annotation outside of plants and animals, leaving a significant gap in understanding less-characterized organisms. Natural variation in terms of signal and noise between organisms and libraries further complicates annotation efforts, particularly for species that may have lost some or all functional RNAi machinery **(Drinnenberg et al., 2009)**. Consequently, these factors create significant challenges for defining locus margins and preventing false annotations. All of these problems are further exacerbated when performing large-scale analyses over many samples and projects, as the risk of exceptional and erroneous samples increases. To address these challenges, this study introduces a data-driven approach that combines threshold optimization and aggressive merging to enhance annotation accuracy. By comparing our methodologies with widely used tools, we demonstrate clear distinctions in how sRNAs are annotated. We show our approach offers robust solutions for producing more consistent annotations across a broader range of organisms, paving the way for improved sRNA characterization in previously underexplored clades.

## Results

### Small RNA data are variable between diverse species

Since the advent of high-throughput sequencing, many species have been studied using sRNA-seq. Searching the NCBI-SRA for “miRNA-seq” libraries reveals tens of thousands of entries (**Fig. 1A**), predominantly from animals, but with a growing representation of plants and fungi counts (**Fig. S2**). However, these libraries are not all equivalent. For instance, library depth varies considerably, with fungi and plants generally exhibiting higher read depths than animals (**Fig. 1B**), possibly due to more recent sequencing efforts having higher depth sequencer-runs. Fungi typically have much smaller genomes than plants and animals, leading to deeper library coverage over genomic space in fungi (10-100x) compared to the other clades (**Fig. 1C**). While this high functional depth (median 1 read per genomic nucleotide) can enhance resolution, it also poses challenges for annotation, particularly for tools optimized for sparsely distributed alignments like those observed in plants.

**Figure 1.**
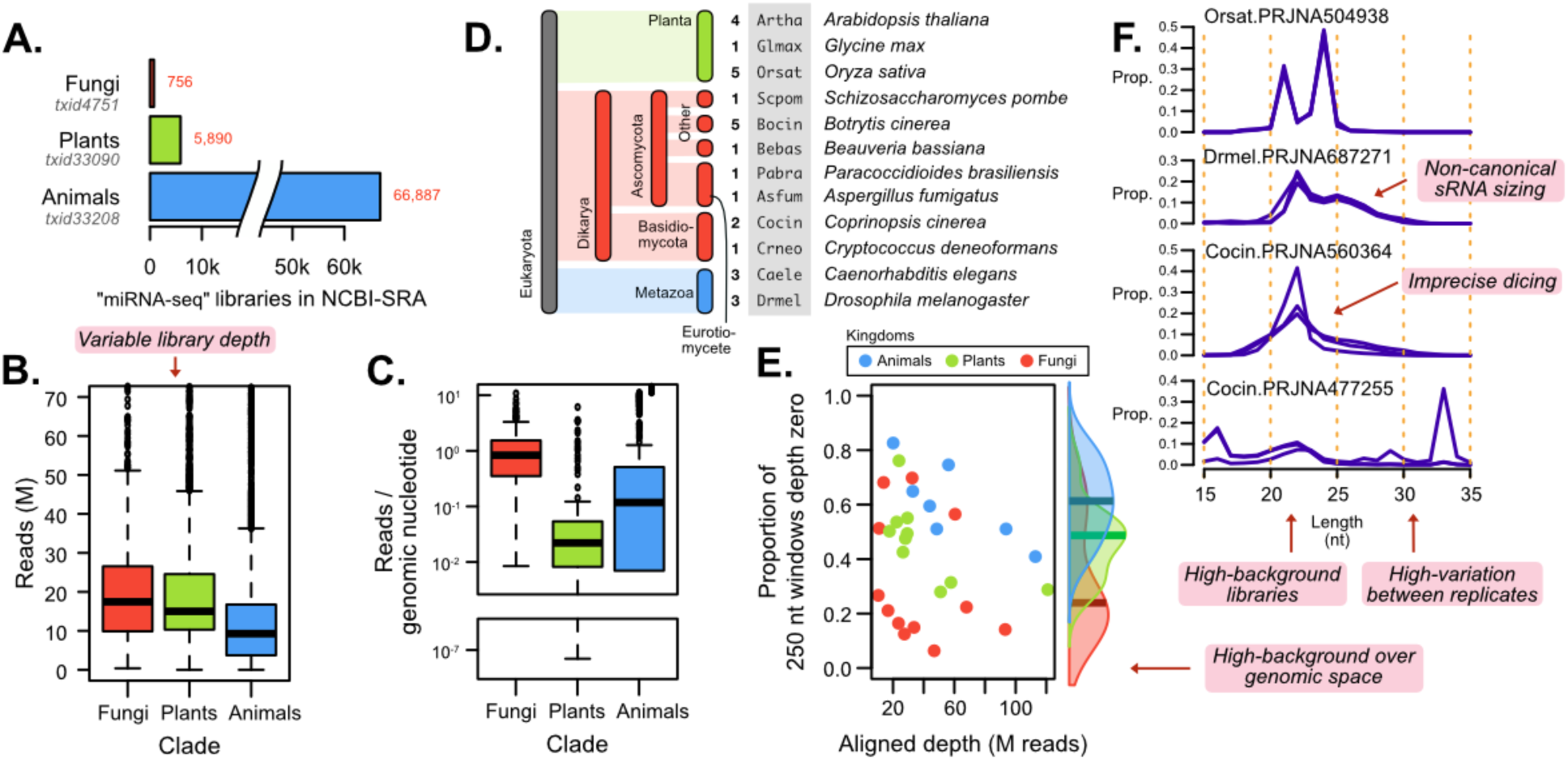
Challenges with sRNA annotation at large. A. Raw counts of “miRNA-seq” libraries in NCBI-SRA. B. Distribution of their raw read-counts and C. their global spatial density over the genome. D. Selection of species included in the test-set of sRNA-seq bioprojects. E. For each bioproject, the proportion of zero-depth 250nt bins plotted against total aligned depth. Points are colored by animals (red), plants (green), and fungi (blue). F. Size-profiles of example alignments of several bioprojects, with replicate libraries shown as overplotted lines.

To explore this relationship in greater detail, we randomly selected a set of publicly available libraries from bioprojects in the NCBI-SRA, including plant (10), animal (6), and fungal (12) projects from several species (**Fig. 1D**). These focused on getting multiple projects for several organisms with variation in sequencing depth, requiring only that they maintained a size-profile that had a distinct sRNA peak (Fig. S3). We aligned wild-type control replicates to their reference genomes and assessed the alignment coverage. By segmenting the genome into 250nt bins, we observed that the fungi have a low-proportion of the genome with no aligned reads (median less than 20%) (**Fig. 1E**). When comparing aligned depth with this zero-proportion, we identified a general trend: deeper libraries exhibit less zero-depth-space (**Fig. 1E**), suggesting that both sequencing depth and genome size contribute to the dense alignments observed in fungi. This ubiquity suggests high noise from regions that are not producing sRNAs or significant transcription.

Looking at some selected size profiles of aligned sRNAs across different species, we observe several challenges associated with real data. For instance, *Orsat* (*Oryza sativa*) represents a characteristic clean profile from a plant, with sharp peaks at 21 and 24 nt (**Fig. 1F**). In contrast, *Drmel* (*Drosophila melanogaster*) exhibits two types of non-coding RNA, 22-nt peak sRNAs and piRNAs around 25-28nt, both of which may not be found by annotators by default. Broader or more diffuse peaks found in some libraries may be an indication of imprecise dicing or other mechanisms specific to an organism (**Fig. 1F**). Library quality presents a major hurdle, as low-quality libraries can severely distort the results of an analysis (*Cocin*.PRJNA477255). Additionally, library depth plays a crucial role in annotation, with large variations in raw reads (Fig 1B) and genomic background (Fig 1E) further complicating the density of reads in the alignment. Finally, discrepancies in library size and profile among replicates justifies careful normalization prior to annotation to prevent disproportionate influence from deep or problematic libraries (**Fig. 1B, F**).

### Annotating sRNAs balancing sensitivity and specificity

Many tools are available for small RNA annotation, varying widely in their scope and strategy (Fig. S1). Here, we focus on annotating sRNAs agnostic of class definitions, leading us to consider ShortStack **(Johnson et al., 2016)** due to its wide usage and command-line interface. Preliminary work with ShortStack version 3 (SS3) revealed difficulty with high-density alignments, such as those observed in fungi. This issue manifested in several locus phenotypes: *over-merging*, where apparently different loci get clustered; *under-merging*, where an apparent larger locus is split into sequential loci; and over-annotation - where large sequential regions of the genome are annotated as loci. We reasoned that these problems stemmed from the way ShortStack3 defines locus margins and proposed that a more refined approach to locus identification, expansion and merging of loci could result in more-realistic alignments.

This motivated the development of our own tool to perform annotations. Here, we address challenges related to alignment density, variation in library size and profile, as well as non-canonical small RNA sizes that may exist in less explored clades. Our approach is implemented within YASMA (Yet another small RNA annotator): a comprehensive software suite for the full-processing, alignment and annotation of sRNA loci (**Fig. S4**). The primary annotation tool for YASMA called “tradeoff” (YTO), follows the procedure outlined in **Fig. 2A** (detailed in Methods and Materials). Briefly, this involves normalizing coverage for each-library individually, balancing influence among replicates. The threshold-depth for annotation is determined using a knee-finding algorithm **(Satopaa et al., 2011)** which balances between annotating the *least genomic space* while annotating the *most reads*. This threshold is then used to define regions which are merged based on proximity and similarity in size-profile, stranding, and expression patterns. Basic filters are employed to make sure that regions have a minimum complexity and abundance to accurately assess their contents (**Fig. 2A**). YTO also allows for library-specific annotation of a sRNA alignment, simplifying the process of alignment, annotation, and analysis for projects with multiple treatments, not all of which should be used for annotation (i.e. pleiotropic mutants).

**Figure 2.**
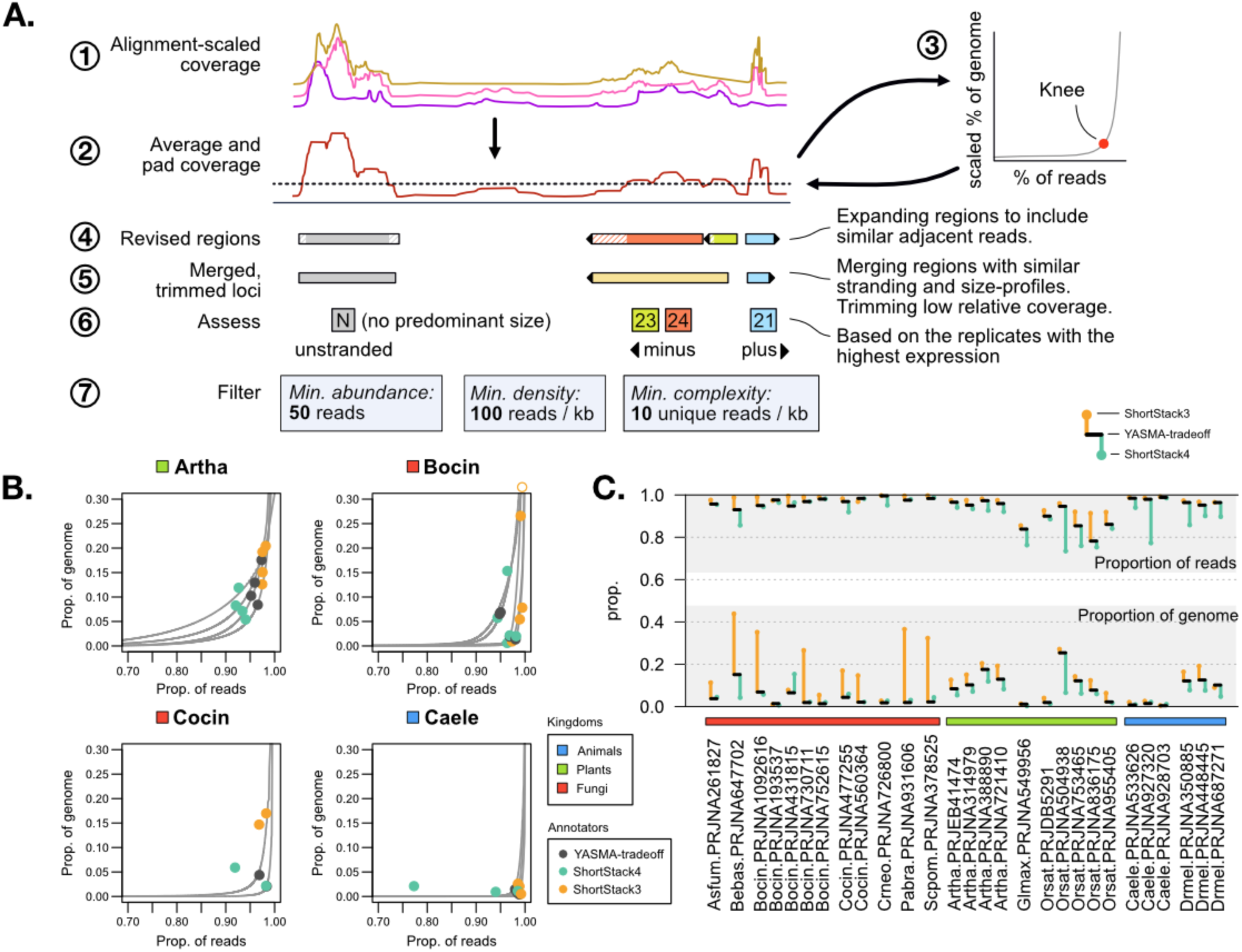
Thresholding sensitivity to avoid over-annotation. A. Overview of the YASMA-tradeoff annotation method, with numbers indicating the progression of steps. B. Example tradeoff curves measured by YASMA-tradeoff for several species (grey), with points showing the selected weighted tradeoff value. Hollow circle indicates a point outside the graphing window. C. The overall annotation rates for all three tools. Proportion of genome included (bottom) and proportion of reads (top) included in annotations are shown, with YTO (black bar), ShortStack3 (point connected to bar by line, orange), and ShortStack4 (same, aquamarine). Alignment depths (M reads) are shown below. Colors represent kingdoms: plants (light green), fungi (bright red), and animals (light blue).

Defining the threshold is one of the critical aspects of annotation by expression. ShortStack version 4 (SS4) is a revision of the ShortStack methodology which adopts normalized-coverage-based cutoff for locus identification, set at 1 read-per-million (RPM). Setting a simple global cutoff in terms of RPM is a simple approach which improves upon the prior strategy, yet this may result in some biases depending on the shape of the alignment.

Comparing this with YTO, we observe similar performance in many organisms, although in some cases SS3 and SS4 produce outlier results (**Fig. 2B**). SS3 tends to annotate much larger portions of the genome across all our test organisms (**Fig. 2C**). This trend is even more pronounced in fungi, where in some species it annotates as much as 40% of the genome as sRNA-loci, echoing the preliminary results. Overall, SS4 and YTO perform similarly (Fig. 2C, S5, S6). The key exceptions are that YTO annotates more sRNAs in plants and animals (Orsat; *Glycine max*, Glmax) whereas SS4 can tend to be more conservative (**Fig. 2B**). A notable point is Orsat.PRJNA504938 (**Fig. 2C**), which has very high depth and a large genome which seemingly produces many sRNAs. YTO and SS3 both define much more of the genome as sRNA-loci, pointing to that this genome produces many loci which are only detectable with high-sequencing depth and might be lost by a strict RPM threshold.

### Locus dimensions and contiguity vary between methods

Contiguity of loci is important for defining discrete, genomic units of sRNA expression. We posited that closely adjacent regions (<500 nt) with similar expression and read profile were likely to be transcriptionally linked.

To assess how loci compare between annotators, we next focused on their attributes. Looking at an example locus in *Coprinopsis cinerea* (*Cocin*, PRJNA477255) (**Fig. 3A**), we see YTO and SS4 are more exclusive in their identification of loci. Here, SS3 appears to exhibit under-merging/over-annotation, with many directly sequential loci. Merging from YTO can be seen with several of the loci, where one YTO locus is found in two or more SS4 loci.

**Figure 3.**
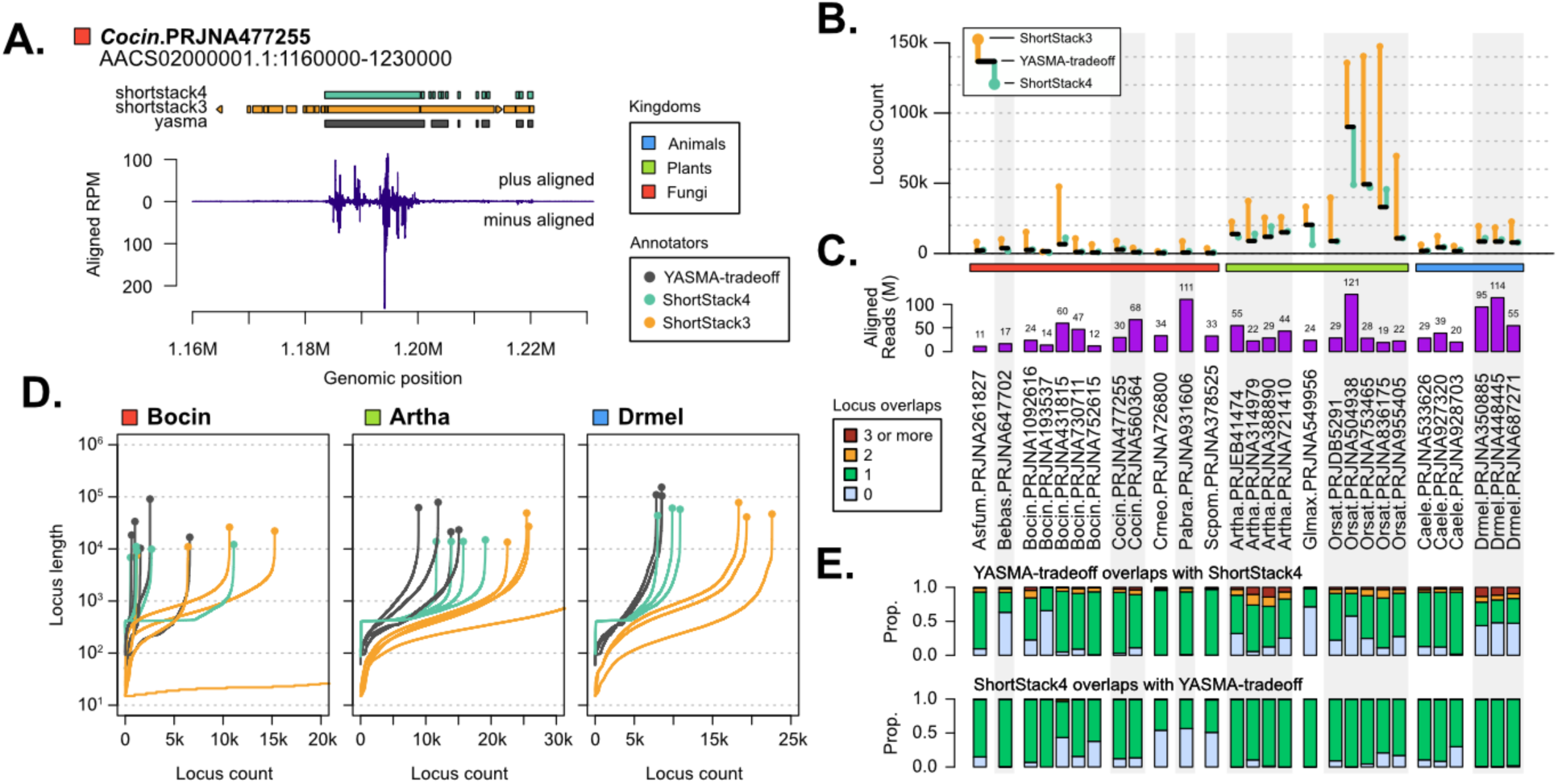
Variable locus contiguity and sensitivity. A. An example of a genomic window, showing annotations for ShortStack3 (orange), 4 (light green) and YASMA-tradeoff (black). Values are shown as coverage in reads-per-million, separating plus-strand (top) and minus-strand (bottom) alignments. B. Annotation dimensions for tested bioprojects, showing the overall locus count and medians for inter-locus gaps and locus lengths. YTO (black bar), ShortStack3 (point connected to bar by line, orange), and ShortStack4 (same, aquamarine). C. Alignment depths (M reads) for each bioproject. D. Locus length plotted against cumulative count of loci for three organisms. Line colors represent the tool as above. Terminal points on lines indicate the largest locus size. E. Counts of the number of ShortStack4 loci which overlap with each YASMA-tradeoff locus (top) and the inverse (bottom). Colors indicate zero (light blue), one (light green), two (orange), and three or more (red) overlapping loci.

This can also be seen by looking at the raw locus counts for all three methods. YTO and SS4 tend to produce similar counts, whereas SS3 consistently identifies a dramatically higher number of loci (**Fig. 3B, S7**). This pattern is not limited to fungi but is observed across virtually all organisms tested. Compared with aligned read-depth (**Fig. 3C**) for these bioprojects, we see again that this rarely is related to locus annotations. The exception is in Orsat.PRJNA504938 where again YTO and SS3 find many more loci than SS4; a possible sign of many unannotated loci due to low normalized depth.

By plotting the cumulative locus count versus locus length, we find that YTO tends to find more longer loci than other methods (**Fig 3D**). In Artha and Drmel, the tools behave predictably between bioprojects in terms of count and length profile (**Fig 3D**). The fungi *Botrytis cinerea* (*Bocin*) presents greater challenges, exhibiting higher variance in locus count for the ShortStack annotations and more stable performance for YTO in fungi. Size profiles for these annotations show SS4 imposes a hard minimum size for loci (around 500 nt), whereas the other tools allow for smaller loci (**Fig 3D**). In general, SS3 finds many more small loci (< 500 nt) than the other methods, supporting the phenotype seen in Fig 3A.

One possible explanation for the identification of fewer and larger loci is that YTO merges multiple distinct regions analogous to those in SS3 and SS4. To explore this, we examined the intersections of loci between tools. This showed that indeed many loci annotated in YTO overlapped with two or more loci identified in the other tools (**Fig. 3D, S8**). This is particularly distinct in SS3, where as much as 40% of YTO loci contain 2 or more SS3 annotations (**Fig. S8**). Looking at the gaps between loci, we confirm that YTO is producing much less closely adjacent loci SS3 and to a lesser degree SS4 (**Fig. S9**). For all tools we find a number of loci in bioprojects which are not found in others. Looking at the expression of these loci in SS3 and SS4, we find most make up less than 2% of aligned reads in total, indicating many loci with very low expression.

### Improved on reducibility in annotations with higher sensitivity

As high-quality annotations are the obvious aim of all these tools, we next tried to assess the confidence of these annotations. Limited validated-data on sRNA-annotations are available in any organism, and are restricted to predominantly miRNAs **(Kozomara and Griffiths-Jones, 2014)**. We decided that reproducibility of annotations between projects could be a good proxy for quality, as loci that are consistently-found between laboratories and experiments are more-likely to be real sites of expression. For those organisms where we tested multiple projects, we performed pairwise intersections of the annotations to measure their similarity. Looking at the overlap of annotated space (**Fig. 4A**, jaccard index), finding that some organisms have highly dissimilar projects and have generally low overlap with all of the tools. We considered that identifying non-annotated regions is also important, especially in high-noise libraries where we might experience more false annotations. For this, we utilized symmetrical comparisons (**Fig. 4A**, simple matching coefficient), which also evaluates the unannotated portions of the genome. Here, we find that YTO and SS4 tend to perform the best, showing that the specificity of these tools yields higher reliability. This is particularly clear in *Bocin* projects where SS3 shows very low reliability.

**Figure 4.**
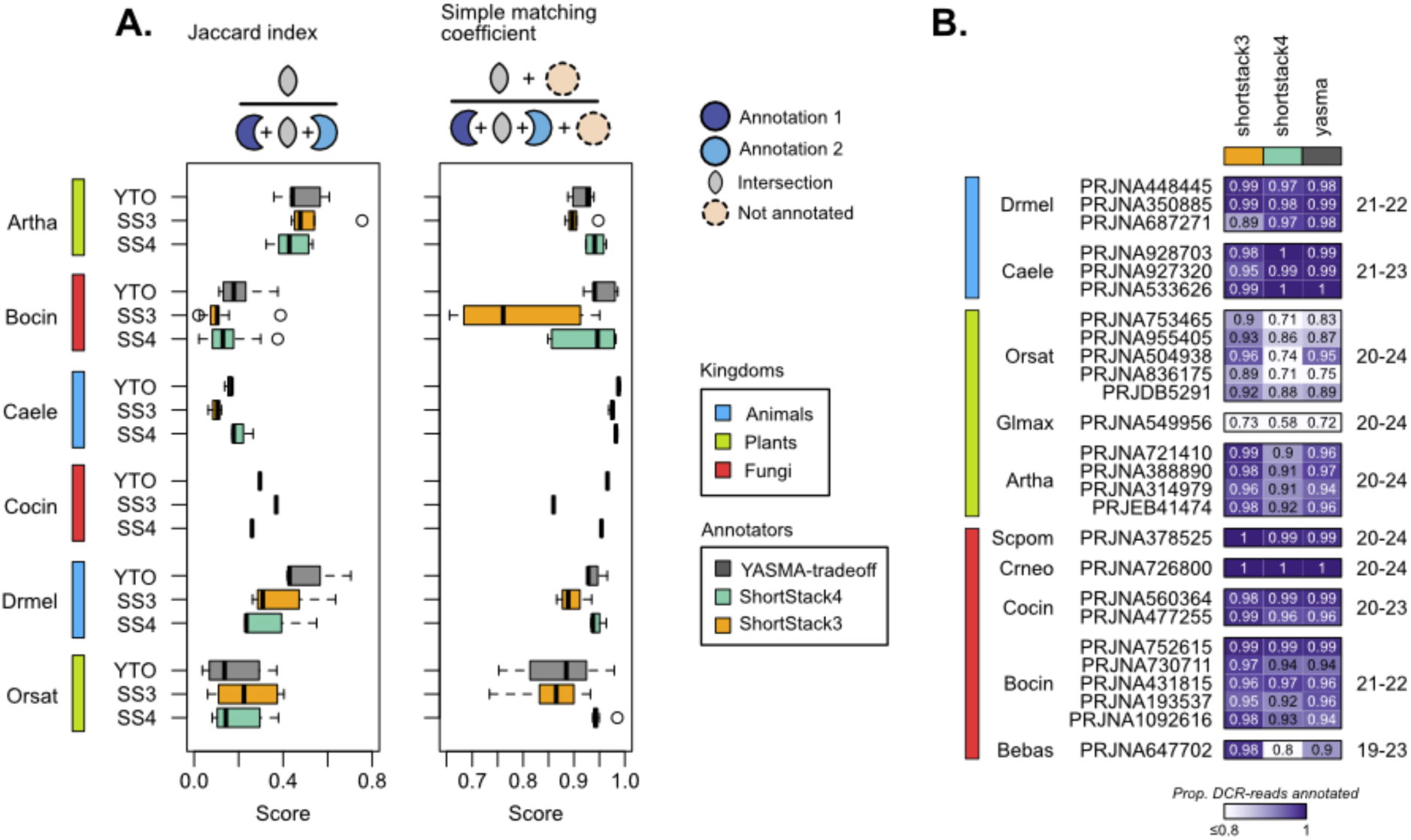
Evaluating annotation reproducibility and sensitivity. A. Pairwise annotation similarity between bioprojects in each organism. Left - jaccard index, right - simple matching coefficient. B. Annotation sensitivity of DCR-derived-reads. DCR-derived size-ranges are shown to the right of the heatmap. Color scale indicates the proportion of Dicer-sized reads annotated.

**Figure 5.**
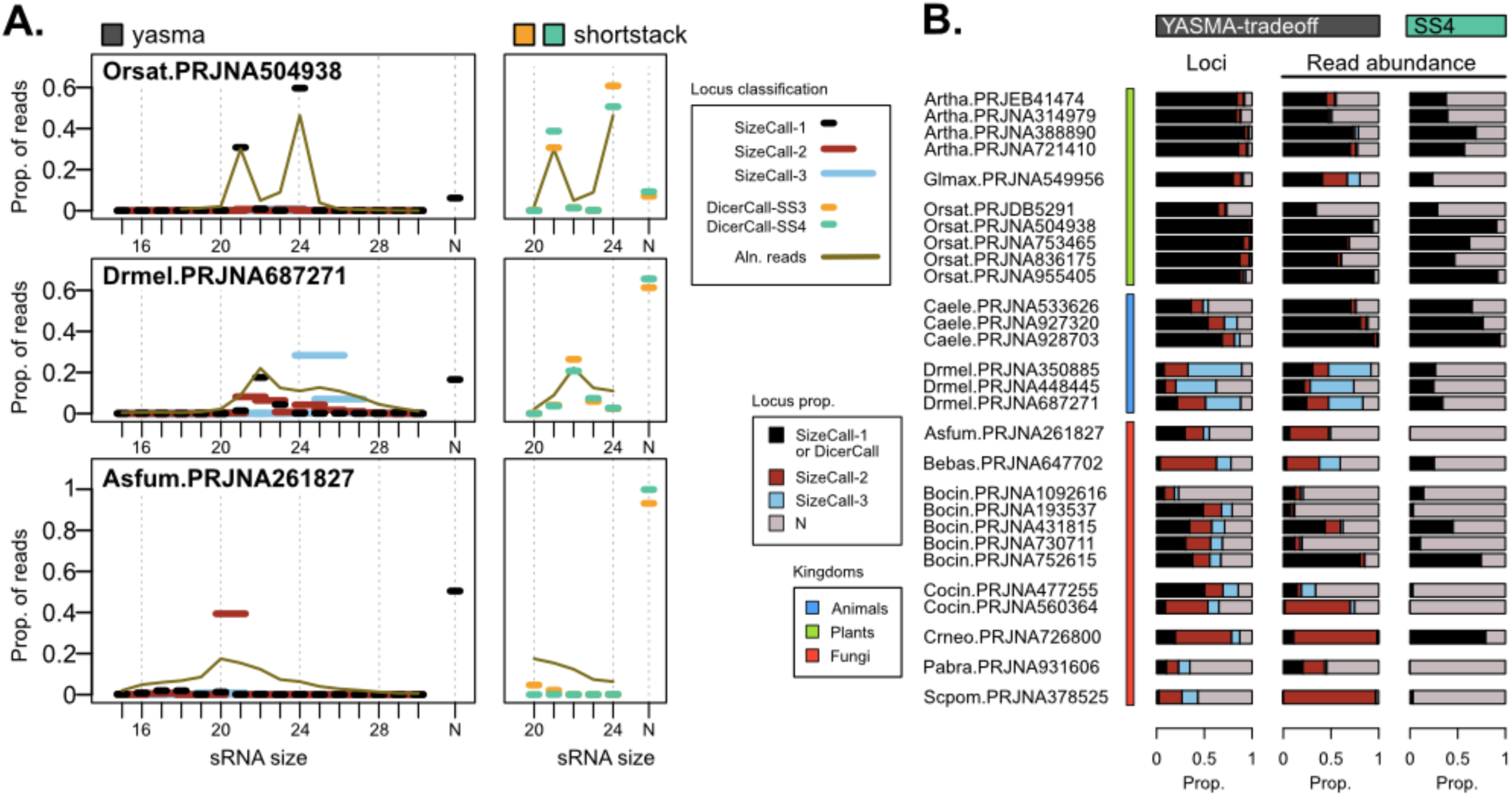
Locus description is methodology-dependent. A. Size profiles of small RNA libraries in several bioprojects. Brown line shows the proportion of reads for each length of read. Lines with circles indicate proportion of all reads that fall into size-categorized loci: “sizecall” for YASMA-tradeoff with locus widths of 1 (black), 2 (dark red), or 3 nt (sky blue), “DicerCall” for shortstack3 and shortstack4. B. Proportion of loci count and abundance for sizecalls-1, −2, and −3 (as described above), and “N” loci (grey). Annotations from YASMA-tradeoff and ShortStack4 (aquamarine) are presented.

**Figure 6.**
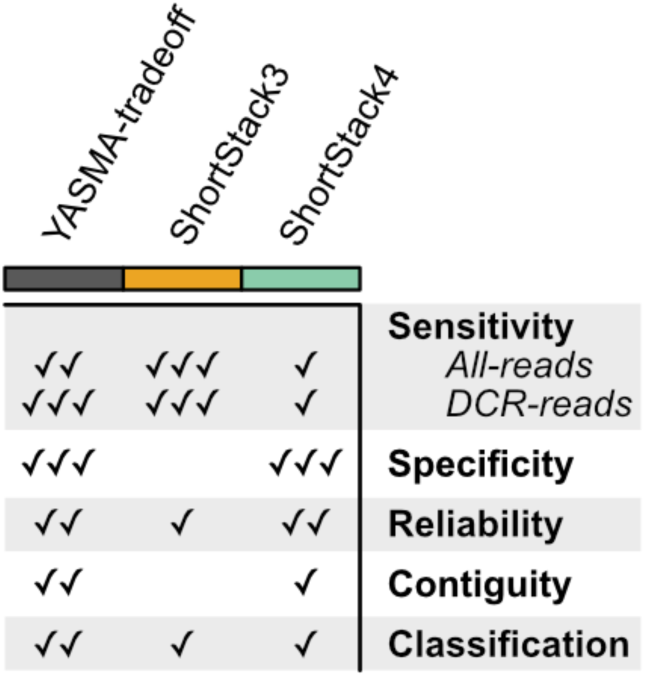
YASMA balances metrics of annotation quality. General summary of annotative quality across several metrics. Relative scores are shown on a scale of 0 to 3 check marks.

We then moved to explore the sensitivity of each of these approaches. Using known-DCR profiles (plants and animals) as well as the size-profiles of our sRNA libraries (fungi) (**Fig. S3**), we built a list of sizes likely to be sRNAs in our samples (**Fig. 4B - right**). By testing the proportion of reads from these size profiles that are in annotations, we can measure the sensitivity of DCR-derived read annotation, considering unannotated reads to be false-positives (**Fig. 4B**). Here, we show that most tools behave similarly in animals and some fungi. In plants, SS4 shows lower sensitivity than the others, most dramatically in those with larger-genomes (*Orsat*, 389 kB; *Glmax*, 978 kB). Overall this points to a higher read-sensitivity in YTO with little cost in reproducibility.

### Capturing imprecisely processed and non-canonical loci

An ongoing concern for annotation in fungi and other yet-unexplored organisms, is that loci may appear different than those described in well-studied systems. The sizes of constituent sRNAs within a locus provide valuable insights into its function, as specific sizes are known to be associated with biological roles and certain machinery/processes. ShortStack infers locus processing using a value called “DicerCall”, which identifies loci where a large proportion of reads fall within a range known to be Dicer-related. Those that pass this test are “called” based on their most abundant size. This is a very powerful tool for inferring a biological role of a locus. However, it also suffers in samples that fall outside of canonical ranges or have higher noise - possibly due to imprecision in processing. YTO adopts a less strict approach, focusing on classifying loci based on the most common contiguous sizes, allowing up to a range of 3 major sRNA sizes. This strategy enhances sensitivity for loci which might have dispersed, but still contiguous sizes, or for loci that are likely not precisely processed by a Dicer protein (e.g. piRNAs). This may be particularly important for exploring sRNA loci in less-characterized organisms, where clearly delineated processing patterns may not exist.

Looking at three example profiles, we can compare how these methods perform by examining the abundance of reads categorized into size-grouping (**Fig. 5A**). In *Orsat*, the results are quite similar between “sizecall” and “DicerCall”, with nearly all reads derived from loci called as 21- and then 24-nt loci. In *Drmel*, 21-nt loci are annotated similarly by both approaches, but reads derived from larger loci within the piRNA range are grouped as “N” in ShortStack. In contrast, YTO classifies these as predominantly 24_25_26 and 25_26_27 loci, greatly reducing the “N” unknown loci. For *Asfum*, we see a highly dispersed sRNA profile centered around 20-nt. ShortStack fails to identify virtually any loci with a defined size-profile, with >95% of reads categorized as “N”. YTO, however, identifies many sRNAs as 20_21 loci, reducing the number of “N”-loci derived reads to approximately 50%.

Looking at the proportions of the “sizecalls” among all test libraries, we can see several that greatly benefit from this approach. Organisms such Bebas (*Beauveria bassiana),* Cocin, Crneo (*Cryptococcus deneoformans*), Scpom (*Schizosaccharomyces pombe*), and Drmel all display large numbers of loci with broader size distributions (**Fig 5B**). Furthermore, for several of these organisms, these larger sized loci represent the majority of annotated sRNA abundance (**Fig 5B**), underscoring the limitations of precise-only analyses and highlighting the value of YASMA’s flexible approach.

## Discussion

### No gold standard for sRNA annotation

This study has been focused on comparing the resulting annotations from YTO and ShortStack tools. A classic approach to this would be binary classification, assessing the annotations in the context of which loci are real/not-real and annotated/not-found. Unfortunately, there remains no gold standard annotation or annotative approach for sRNA-producing loci. Several annotations do exist in databases **(Kozomara and Griffiths-Jones, 2011)** and repositories of annotations **(Lunardon et al., 2020; Cardoso et al., 2018)**. But, using these as a standard to compare annotations is inappropriate, as their own annotations are subject to the methodology-used (Fig. S1). We instead focus on indicators of quality based on comparing our annotations and general characteristics. We have proxies for some key metrics: proportion of reads annotated is similar to sensitivity and proportion of genome annotated is similar to specificity (Fig 2). Additionally, we compared annotation reliability (Fig 4), DCR-derived-read sensitivity (Fig 4), and annotation over/under-merging (Fig 3, S9).

Among these metrics, different approaches have different strengths, outlined generally in Fig 6. SS3 shows extreme sensitivity, annotating the highest percentages of reads. This comes at the cost of annotating vastly more loci than the other tools, many of which are directly adjacent to other loci and small. It also struggles in some organisms, namely fungi, annotating large regions of the genome as single loci. SS4 is more conservative, producing lower counts of more isolated loci, which are in-turn more reliable. This has the cost of lower sensitivity than the other methods, particularly in plants. YTO manages to balance the benefits and problems of these approaches. It retains high read-annotation sensitivity, matches SS4 in terms of reliability, and produces the most-contiguous loci of any tool.

### Classifying loci is key for understanding role

Noise plays a major factor in sRNA annotation with high-depth libraries, as separating sRNA loci from the noise can be a challenge. RNAs in the size-range of sRNAs can be produced as the natural product of RNA-degradation **(Houseley and Tollervey, 2009)**. Deciphering whether an expressed region is derived from directed processing, often by a DCR protein, is key to identifying loci that may have a regulatory role. DCRs produce specific-sized sRNAs **(Henderson et al., 2006)**, whereas degradation produces RNAs with irregular or random sizes. Similarly, specific AGO proteins preferentially bind to particular sRNA lengths **(Mallory and Vaucheret, 2010)**, directly linking the size of an sRNA locus with its biological function. Here, ShortStack takes a conservative approach, while applying a standard realistic for the precision found in plants. In contrast, YTO adopts a broader approach, allowing for many more loci to be categorized and assessed based on their observed characteristics. While this approach weakens the ability to determine whether locus is DCR-derived, it opens up possibilities for analyzing other small RNAs, particularly in cases where DCRs act differently, or cryptic dicing might occur **(Yang et al., 2016; Lee et al., 2010)**. In these cases, strong genetic evidence is required to confirm which DCR performs specific processing, meaning that any predictive tool is likely not sufficient.

### Aggressive merging allows for more realistic loci

Locus merging is an important aspect of sRNA annotation. Transcription of sRNAs varies between classes, with some derived from processed POLII transcripts (miRNAs, tasiRNAs, phasiRNAs) **(Zhan and Meyers, 2023)** and others likely from many one-to-one transcription events (siRNAs) **(Blevins et al., 2015)**. These make annotation contiguity challenging, as there can be large gaps in coverage between regions that are likely transcriptionally linked. YTO makes a strong effort to merge highly similar loci at long distances (500 nt), forming much more contiguous annotations. This approach also impacts downstream techniques, as inflating one locus to multiples can affect the sensitivity of techniques like differential expression analysis.

Faulty annotations represent a major challenge for biological interpretation. Loci which are arbitrarily small, large, or divided can greatly alter the interpretation of what they are and what they do. This is evident in miRNA, where low-confidence annotations **(Kozomara and Griffiths-Jones, 2014; Axtell and Meyers, 2018)** have influenced many subsequent publications. Annotators that produce higher rates of false annotations for sRNA loci run the risk of propagating false interpretations of classes, frequency of loci, and ultimately function of a given sRNA. By imposing stricter standards as YTO and SS4 show, this greatly reduces these risks.

### Reliable methodologies are essential for new frontiers

With the rapid expansion of sequencing technology and data, an increasing number of projects aim to uncover meta-level trends within the genomes of organisms. This also includes the field of sRNAs, where numerous repositories of data have been compiled, including kingdom-wide studies **(Lunardon et al., 2020)**, sRNA class-wide datasets **(Guo et al., 2022)**, and even pan-target analyses **(Huang et al., 2022; Liu et al., 2021)**. All these approaches leverage consistent methodologies for large-scale analyses, allowing for powerful comparisons across conditions, species, classes of sRNA and more. We also show that YTO is well-suited to the many classes of sRNAs, including those with non-canonical sRNA sizing, pushing back on these barriers. Other possible applications of this tool could be with other types of non-coding RNA, including those derived from other RNA sources (tRNA, rRNA) **(Martinez et al., 2017; Cherlin et al., 2020)** or alternative pathways (bacterial sRNAs) **(Li et al., 2013)**. Further extending this approach with high-throughput methods of identifying targeting and function **(Yan et al., 2024; Horlacher et al., 2023)** promises to link these annotations to their biological role.

Considering the large quantity of publicly available sequencing data, repository approaches linked to reliable methodologies show great promise **(Lunardon et al., 2020)**. We can leverage replication within a project and between projects, with the idea that consistent annotations are inherently more robust and likely to be real. This also opens the door for evolutionary studies, looking for conservation of locus type and specific loci on a clade-wide scale. This is particularly relevant for underexplored clades, where few works have shown signs of conservation **(Johnson et al., 2022)** and a lack of common annotative approach have proved to be a serious hurdle.

## Conclusion

Here, we present a comparison of how different approaches perform in sRNA-annotation, demonstrating that coverage-normalized methods like SS4 and YTO build much more consistent annotations. YTO uses threshold identification and merging to annotate a maximal amount of reads while retaining specificity. YTO’s merging constructs highly contiguous loci with more sensitive descriptions of size profiles, enabling a broad perspective of sRNAs. These annotators effectively address major challenges in terms of clade-wide analyses, which involve significant variation in sRNA libraries (quality, size, depth), genomes (sRNA types, expression, sRNA machinery), and functions. Here we show an important advancement in methodologies addressing sRNA annotation, allowing for greater insights into what sRNAs are made in organisms spanning the whole tree of life.

## Methods and Materials

### Assessing and gathering public data

To find global metrics of public sRNA-seq data, we searched the NCBI using edirect tools. Our term looked for libraries matching the strategy “miRNA-seq” and one of the TaxIDs for fungi (txid4751), plants (txid33090) and animals (txid33208). Reference genomes were sourced from the NCBI using the datasets tool, using metadata to calculate global metrics. We focused our strategy on a small selection of samples, including several species with 3 or more distinct projects (*Bocin, Artha, Drmel, Orsat, Caele*). All bioprojects and libraries associated with this project are shown in Table S1. Phylogenetic classifications between sample organisms are derived from NCBI taxonomy.

Assessing sRNA annotator citations was done using NCBI esearch in the pubmed database for publications related to the small RNA annotation tool. Citation counts were compiled from esearch, combining related publications into families (Table S2).

### Processing and aligning libraries

YASMA was used for managing all processing and alignment steps for the test libraries. This performs a sRNA-optimized strategy for processing steps from one primary directory, using the following tools and steps. Libraries were downloaded using prefetch and fasterq-dump through the NCBI SRA toolkit (YASMA-download). YASMA-adapter was used to identify common 3’ adapter sequences found in the libraries, which were then trimmed using cutadapt **(Martin, 2011)** with the following settings ‘cutadapt - a [3p_adapter_sequence] --minimum-length 15 --maximum-length 50 -O 4 --max-n 0 --trimmed-only [fasta]’. Trimmed libraries were aligned to the reference genome (Table S3) using YASMA-align, which uses bowtie1 **(Langmead et al., 2009)** and follows the unique-weighting strategy for multi-mapping reads used in ShortStack3/4 and described in **(Johnson et al., 2016)**. In short, this aligns reads with up to one mismatch, and places multi-mappers with a weighted-random selection based on the number of nearby uniquely mapping reads.

### Small RNA annotations

Annotations were done using several approaches. Annotations for ShortStack3 and 4 were done using the same alignments as above, using versions 3.8.5 and 4.0.2. Annotations using YASMA-tradeoff are described here in detail.

YTO first reads the alignment encoding and calculates coverage for each library separately by summing all reads in a sliding user defined window (default: 250 nucleotides). Padding is done to expand region boundaries, again using sliding windows instead calculating ‘max()’ (default: 100 nucleotides). Coverages are normalized to reads-per-aligned-billion reads (RPB). Positional coverages are then averaged between replicates by median and between experimental conditions by mean. Condition averaging is not applicable in this work as only a single condition was used for the test bioprojects. Annotation read- and genome-percentage tradeoffs are then calculated based on the average coverages. The tradeoff value (in RPB) is identified using the approach outlined in kneedle **(Satopaa et al., 2011)**, generally balancing more reads with less genomic space.

YTO then identifies candidate regions based on positions which exceed the RPB threshold. Regions are expanded and/or trimmed to define the boundary where expression is largely lost (<5% of the expression at the edge). Regions are merged sequentially if they meet all the following standards: 1) they must have overlapping size profiles or have no-predominant size, 2) they must have similar a proportion of reads that come from the top strand (a difference in proportion no greater than 0.5 by default), and 3) they must be close enough (less than 500 nucleotides by default). Loci are merged greedily by this method, with recalculation for the bridge area between merged regions.

Loci are finally filtered based on three criteria: 1) minimum abundance (50 reads), 2) minimum abundance density (100 reads / kBase), and 3) minimum complexity (10 unique reads / kBase). These ensure loci are deep and complex enough to be usefully analyzed.

### Genomic intersections

Intersections and comparisons of loci were performed using the GenomicRanges package in R **(Lawrence et al., 2013)**. Jaccard index was calculated between annotations using the following formula: 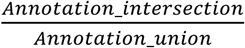. Simple matching coefficients were calculated using the following formula: 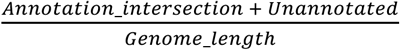.

## Data and code availability

All sequencing data for this study are available in the NCBI-SRA and are outlined in Table S2. All genomes used for alignments are from the NCBI genome database and are enumerated in Table S3. The YASMA annotation suite is available at https://github.com/NateyJay/YASMA. Installation is performed by simply running the yasma.py script from the github repository folder, with more explicit instructions available on its page. YASMA has been tested in Mac-OS and linux systems.

## Author contributions

NRJ conceived of the project, and designed and implemented YASMA. NRJ and FGT performed test annotations, analysis, and troubleshooting of the YASMA methodology. BBG performed the analysis of available annotators. FGT and BBG searched and annotated the metadata of test projects and libraries.

## Supplemental materials

**Figure S1.**
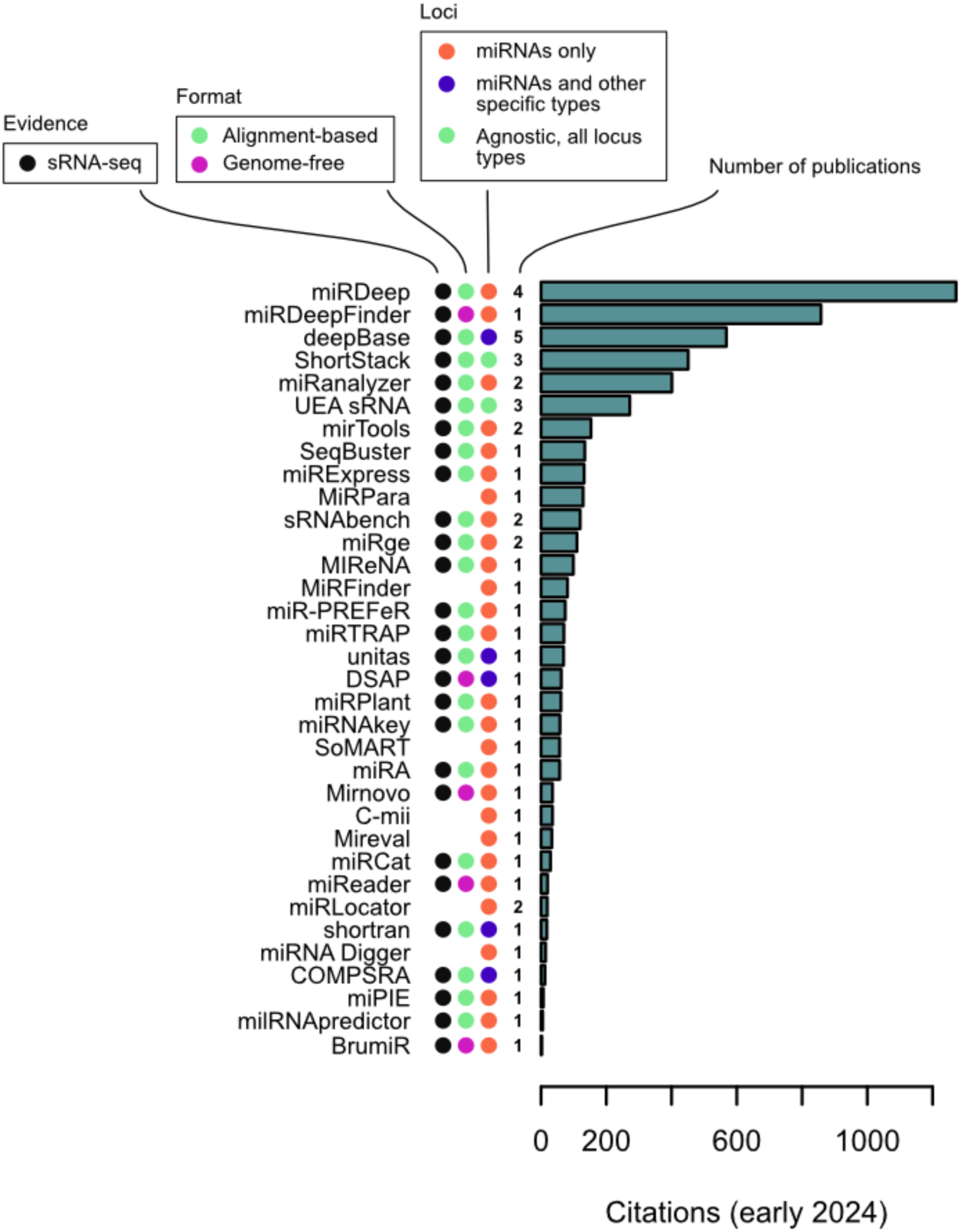
annotator overview. Overview of published sRNA annotators. Tools with multiple publications and versions are aggregated by citation count, listing the number of publications containing this information (Table S1). Tools are described in terms of their scope and strategy, first identifying which are sRNA-seq based. Format of annotation and what locus-types are described are also shown with the described colors.

**Figure S2.**
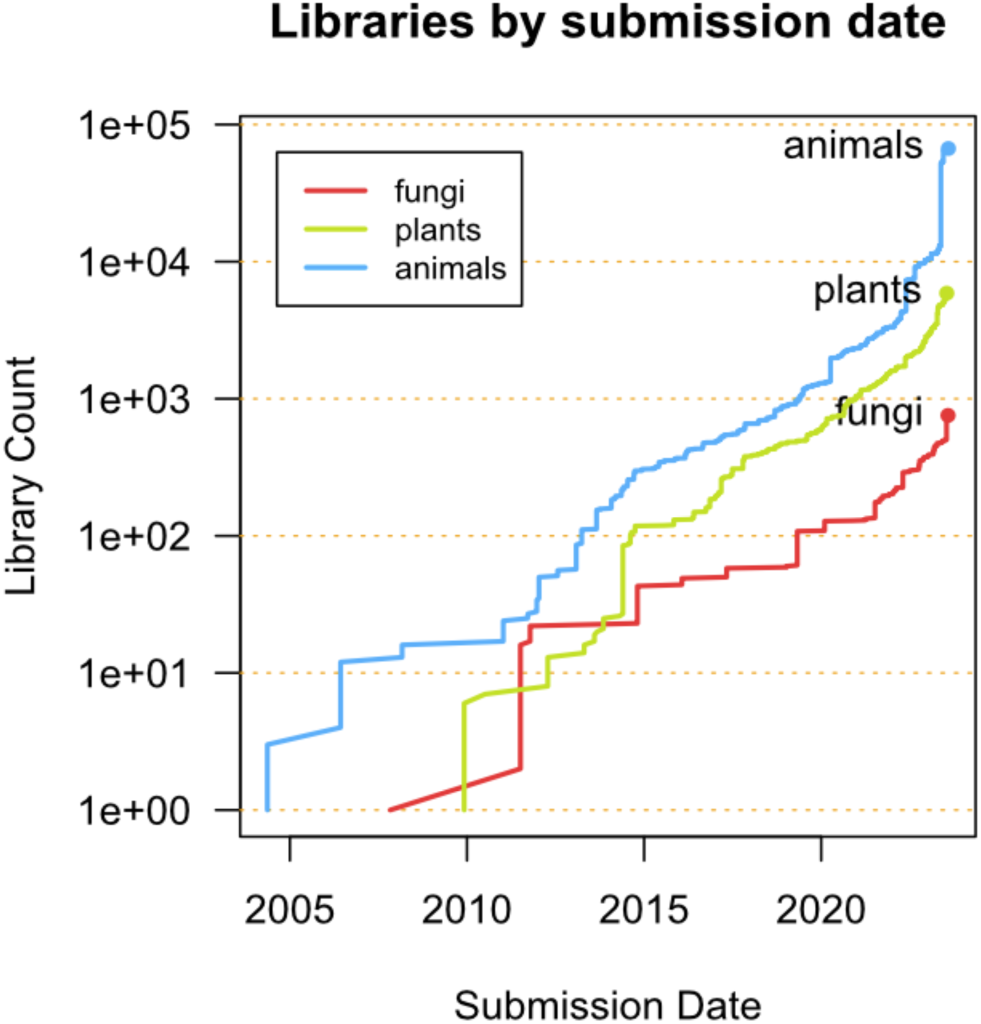
NCBI assessment. Cumulative count of “miRNA-seq” libraries in the NCBI-SRA, aggregated by sample kingdom.

**Figure S3.**
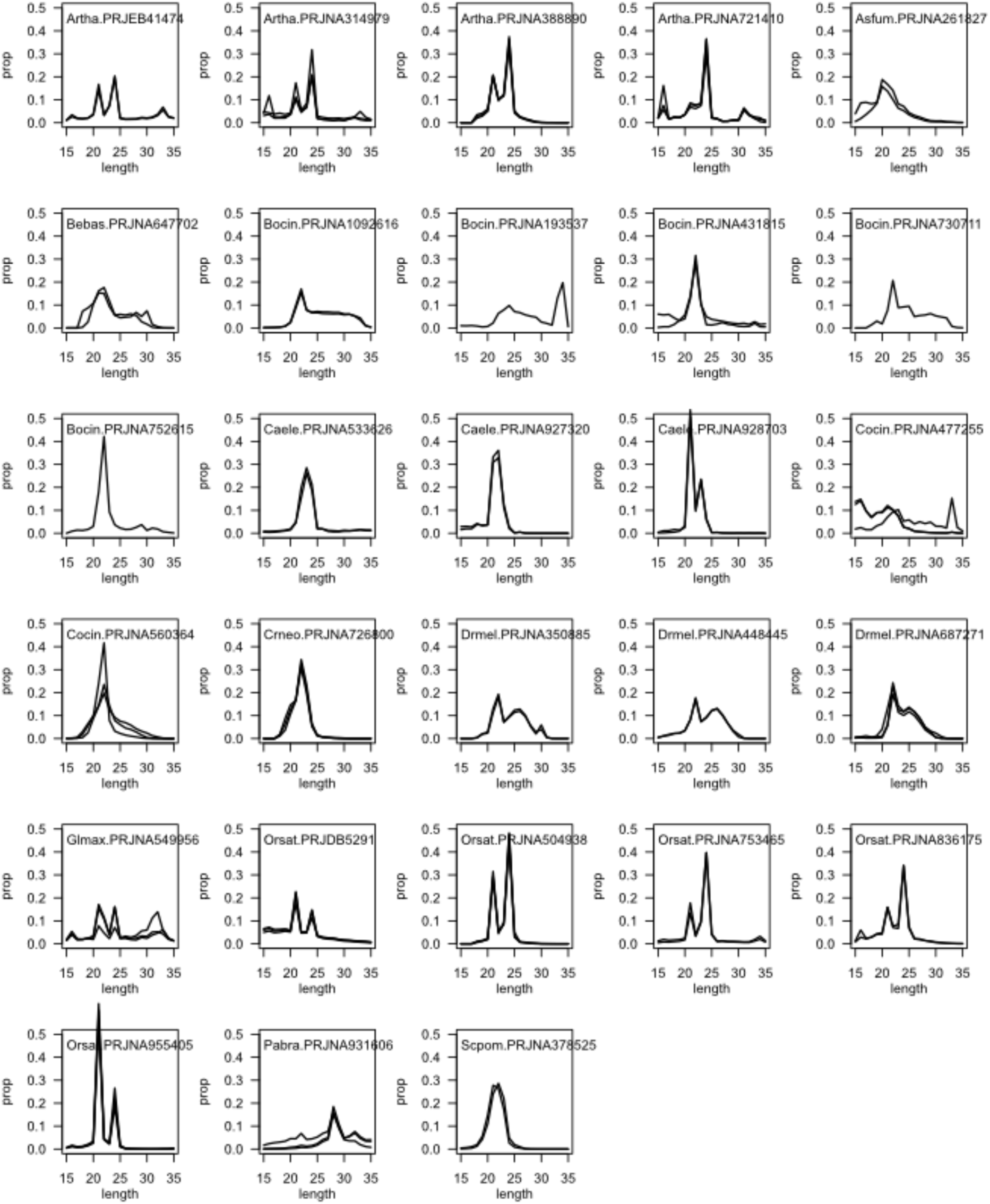
Size profiles of aligned sRNAs. Size profile of all projects included in this analysis. Profiles are proportions of all aligned reads, with libraries shown as different overplotted lines.

**Figure S4.**
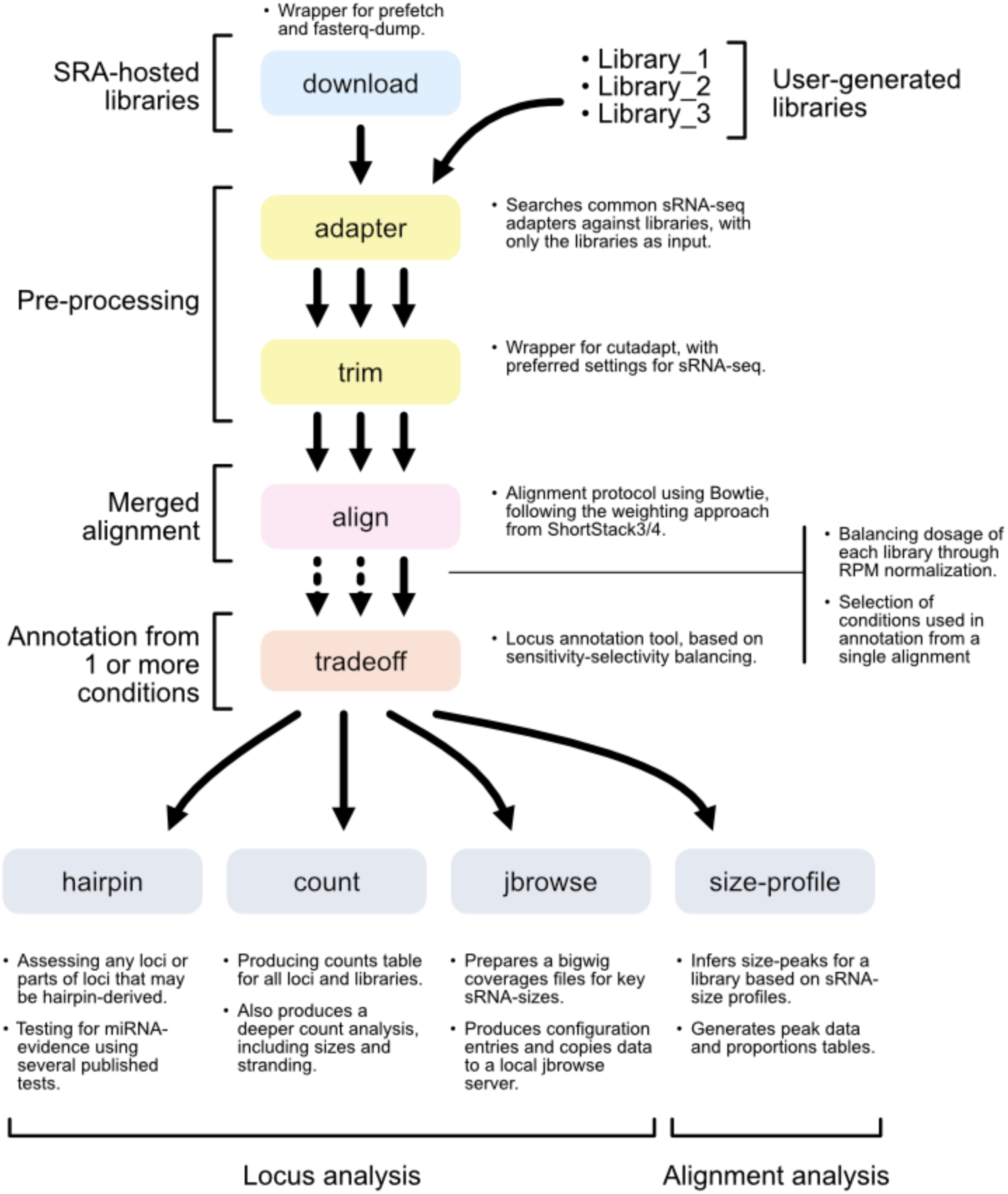
Overview of YASMA. Block diagram indicating the modules and general analysis pipeline for YASMA. Tradeoff is the subject of this work.

**Figure S5.**
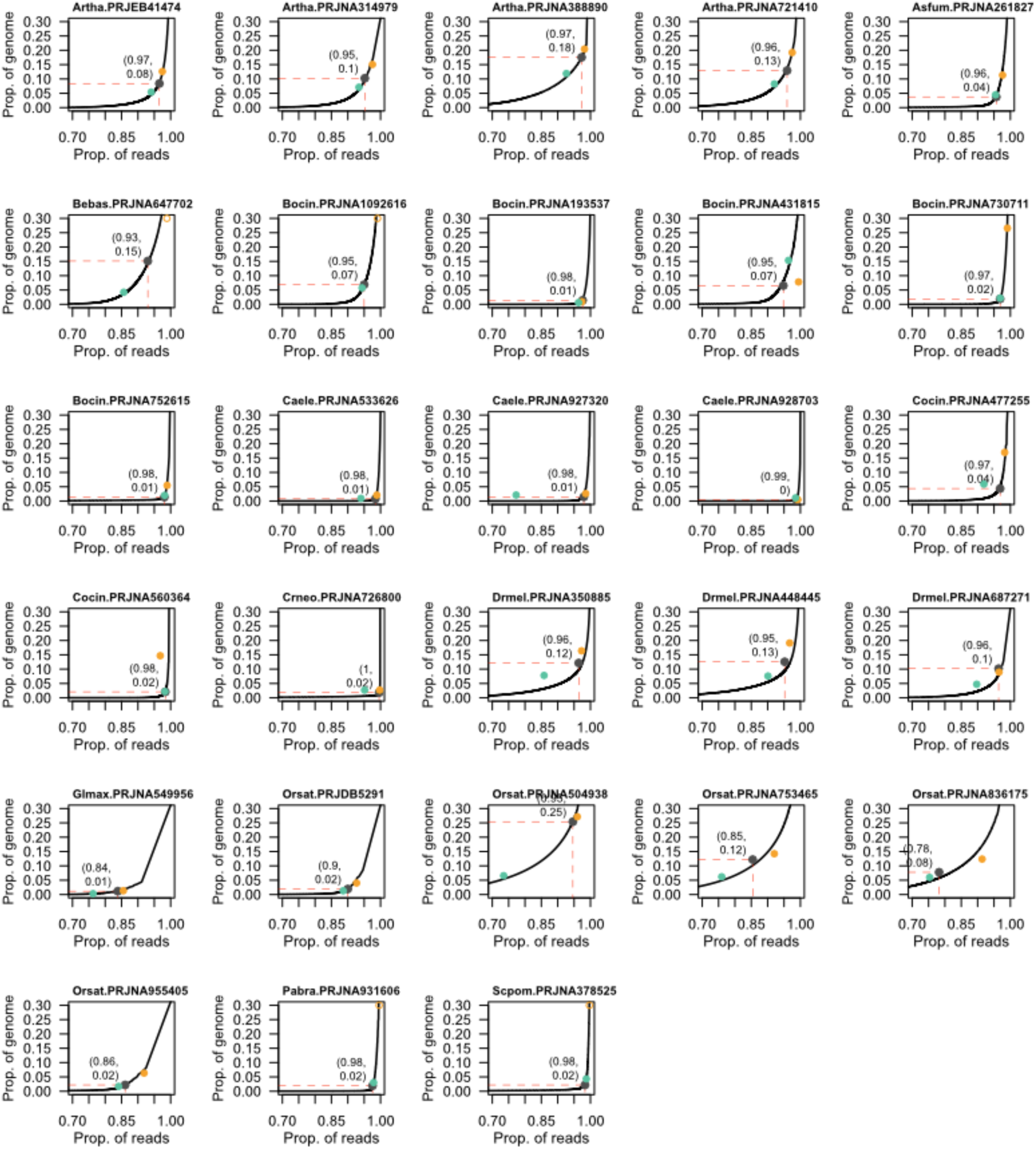
tradeoff curves. Tradeoff curves of the proportion of genome annotated vs the proportion of reads annotated for each project. Curves are calculated by YASMA-tradeoff, with the final annotation rates reported for each tool: YASMA-tradeoff (black), ShortStack3 (orange), and ShortStack4 (aquamarine).

**Figure S6.**
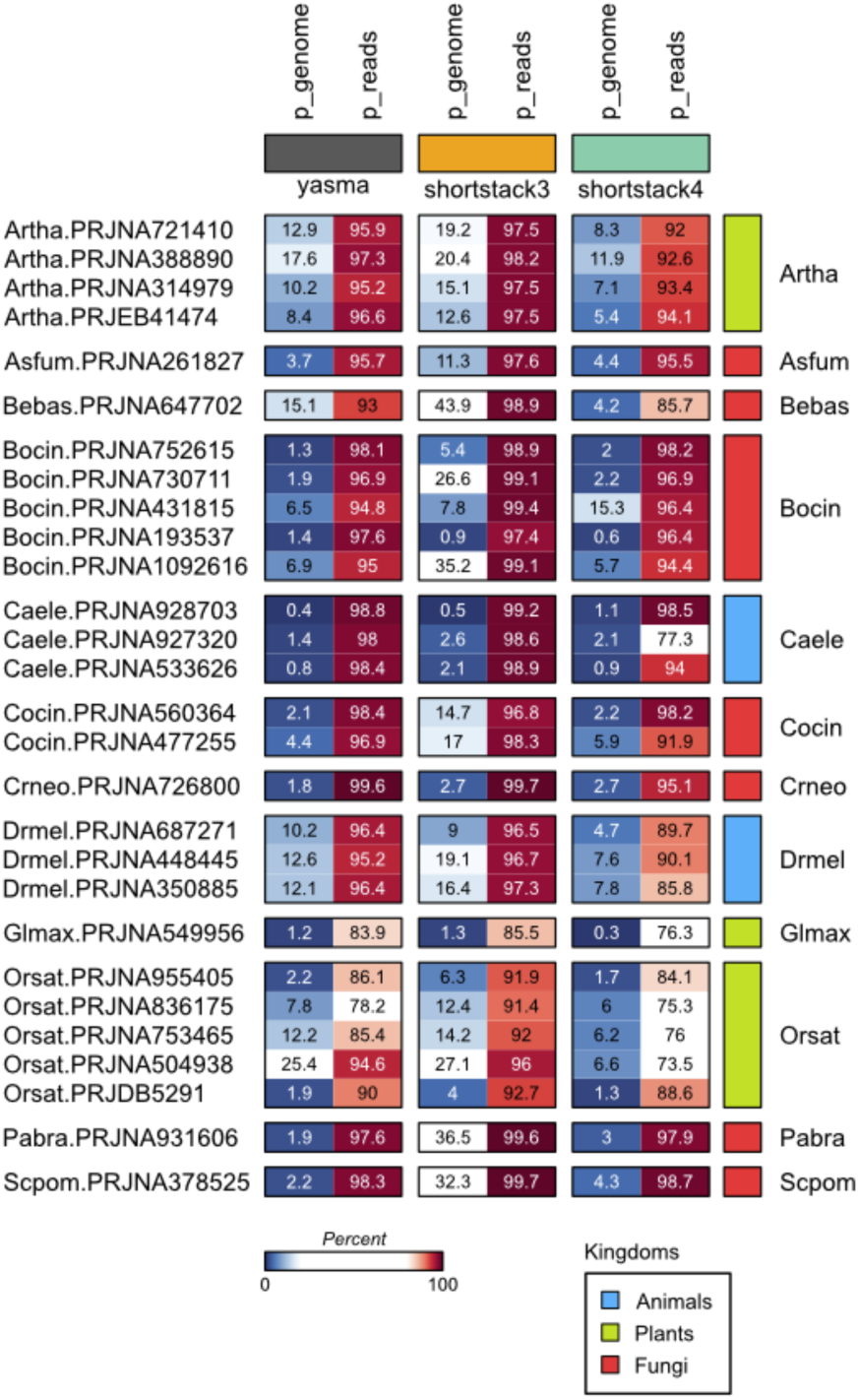
annotation rates for all tools and projects. A summarization of figures 2C and S5, showing the read and genome annotation rates for each annotation. Kingdoms are identified by colors: animals (light blue), plants (light green), and fungi (bright red). Percents are shown as color scales, with blues shown from 0-20%, white for 20-80%, and reds for 80-100%.

**Figure S7.**
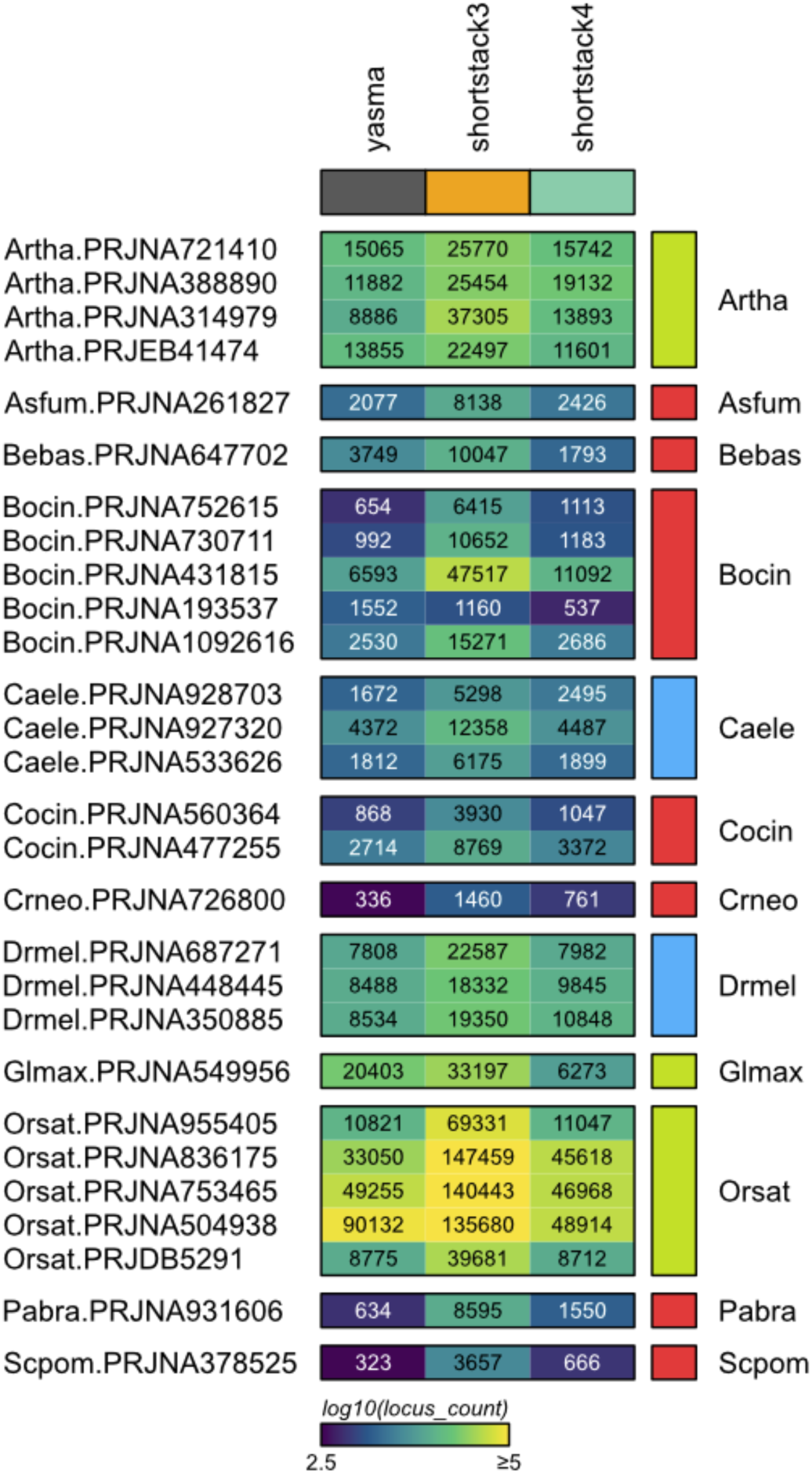
locus counts. Heatmap of locus counts for each project and annotator. Kingdoms are colored on the right as plants (light green), animals (light blue), and fungi (bright red).

**Figure S8.**
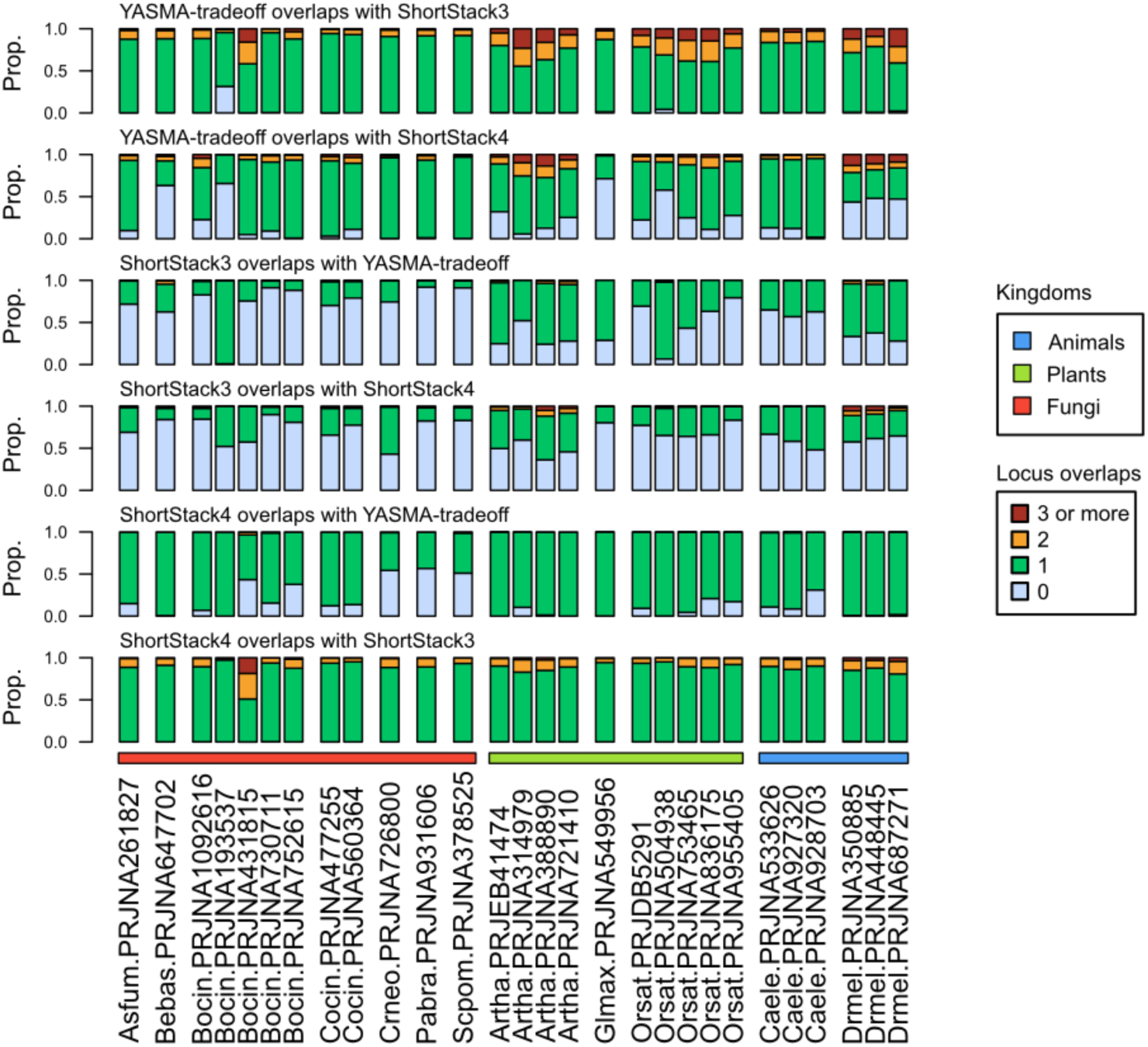
locus overlap between annotations. An extension of figure 3E, showing pairwise overlaps for all annotations.

**Figure S9.**
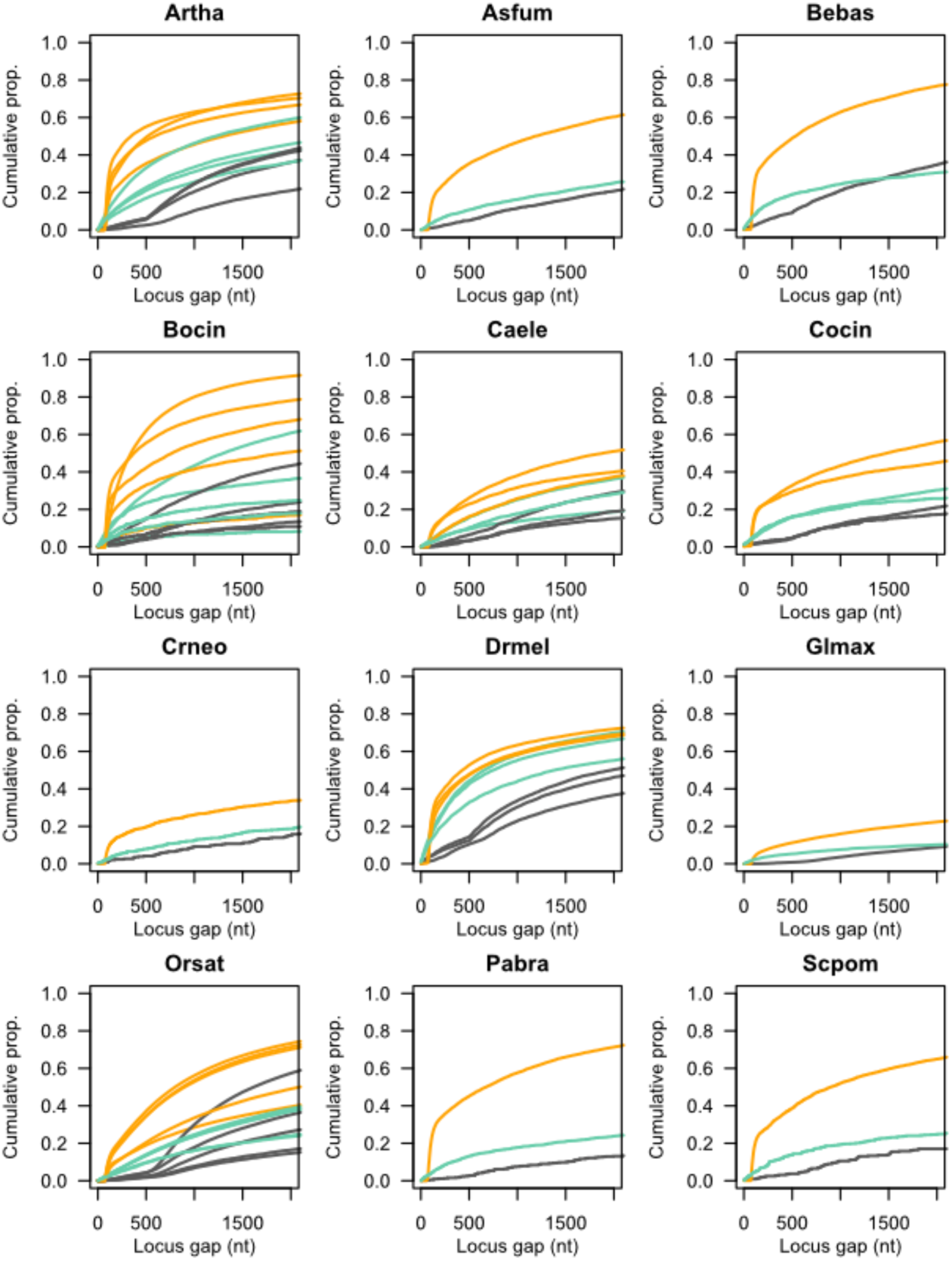
Inter-locus gaps for annotations. Inter-locus gap distances are represented as a cumulative density function for each annotator, aggregated by organism. Lines represent specific annotations and line color indicates the annotator: YASMA-tradeoff (black), ShortStack3 (orange), and ShortStack4 (aquamarine).

*Table S1 - Table of tool publications*

*Table S2 - Table of libraries*

*Table S3 - Table of genomes*

